# Transposable elements drive the evolution of genome streamlining

**DOI:** 10.1101/2021.05.29.446280

**Authors:** Bram van Dijk, Frederic Bertels, Lianne Stolk, Nobuto Takeuchi, Paul B. Rainey

## Abstract

Eukaryotes and prokaryotes have distinct genome architectures, with marked differences in genome size, the ratio of coding/non-coding DNA, and the abundance of transposable elements (TEs). As TEs replicate independently of their hosts, the proliferation of TEs is thought to have driven genome expansion in eukaryotes. However, prokaryotes also have TEs in intergenic spaces, so why do prokaryotes have small, streamlined genomes? Using an *in silico* model describing the genomes of single-celled asexual organisms that co-evolve with TEs, we show that TEs acquired from the environment by horizontal gene transfer can drive the evolution of genome streamlining. The process depends on local interactions and is underpinned by rock-paper-scissor dynamics in which populations of cells with streamlined genomes beat TEs, which beat non-streamlined genomes, in continuous and repeating cycles. Streamlining is maladaptive to individual cells, but delivers lineage-level benefits. Streamlining does not evolve in sexually reproducing populations because recombination partially frees TEs from the deleterious effects they cause.

## Introduction

Prokaryotes and eukaryotes have distinct genome architectures. In general, prokaryotes have small streamlined genomes where up to up to 90% of DNA is host-essential (Koonin & Wolf 2008; Silby et al. 2009). In contrast, eukaryotes have large genomes (Gregory 2001; Oliver et al. 2007) with only a small proportion encoding host-essential proteins (Taft et al. 2007). The intergenic space of eukaryotes is populated by numerous repetitive sequences (Jelinek & Schmid 1982; Kidwell 2002; Jurka et al. 2007), many of which are transposable elements (TEs) or remnants thereof. As TEs can replicate independently of hosts, proliferation of TEs is thought to have driven genome expansion in eukaryotes (Kidwell 2002; Gregory 2005; de Albuquerque et al. 2020). Prokaryotes, however, also harbour TEs in intergenic spaces and yet have streamlined genomes (Stern et al. 1984; Lupski & Weinstock 1992; Siguier et al. 2006; Treangen et al. 2009; Bertels & Rainey 2011). If TEs play a role in determining the large genomes of eukaryotes, then why are bacterial genomes more streamlined?

To understand the relationship between TEs and genome architecture, it is necessary to consider mechanisms underpinning TE persistence. In asexual organisms the long-term fate of TEs within a given lineage is extinction (Park et al. 2021). Opportunity for long-term persistence depends critically on ability to periodically invade new lineages via horizontal gene transfer (HGT) (Doolittle & Sapienza 1980; van Dijk et al. 2020). In eukaryotic populations, TEs are maintained by sex (Wright & Finnegan 2001; Kent et al. 2017).

Although TEs can become linked to ecologically relevant genes and thus confer direct benefits to their hosts, the majority incur measurable fitness costs (Startek et al. 2013; Consuegra et al. 2021). Marginal costs stem from need to replicate the additional DNA that is generated by TE duplication, but more substantive costs arise when TEs integrate into – and inactivate – host-essential genes. Assuming TEs insert at random, then the risk of gene inactivation is directly related to the proportion of host-essential DNA. In other words, TE infection is likely to be costly for bacteria with streamlined genomes, which have a high proportion of host-essential DNA, and less costly for eukaryotes, which harbour large stretches of non-coding DNA.

Here, we present a co-evolutionary model of TEs and their host genomes. The model explicitly considers hosts with a genome that contains stretches of coding DNA and non-coding DNA. Host genomes can be infected by TEs via uptake of extracellular DNA (eDNA). Once infected, TEs replicate within genomes, where integration into essential genes results in cell death and lysis. Lysis liberates TEs back into the eDNA pool. Our model shows that TEs can drive the evolution of genome streamlining in asexual organisms. The process depends on local interactions and is underpinned by rock-paper-scissor dynamics in which populations of cells with streamlined genomes beat TEs, which beat non-streamlined genomes, in continuous and repeating cycles. Genome streamlining does not evolve in a sexually reproducing populations because recombination partially unlinks TEs from the deleterious effects they cause. Together, our findings provide support for a previously unrecognised role of TEs in the evolution of genome streamlining.

## Results

### An *in silico* model of the co-evolution of TEs and their host genomes

To understand how the interaction between TEs and cells shapes genome architecture, we present an individual-based model of co-evolving transposable elements (TEs) and host genomes packaged within cells. We first focus on simple bacterial-like cells, which engage in horizontal gene transfer (HGT) via environmental pools of DNA, but later extend the model to encompass sexual reproduction. A brief overview of the model is given below. A full description can be found in the methods section.

Individuals are simple cells that carry a genome with three distinct genetic elements: **i)** ten host-essential genes (type A-J) which are necessary for survival/reproduction of the host, **ii)** TEs, which are slightly costly to the host, and **iii)** non-coding DNA which provides no function but also carries no cost (**Figure 1a**). Elements are represented as a linear sequence like “pearls on a string” (Hogeweg et al. 2016) and can be exchanged and recombined through different mutational processes (see **Figure 1b**). For example, single gene duplications may result in redundant gene copies, subsequent gene inactivation may result in the generation of non-coding DNA, and further deletions/duplications may expand or reduce the amount of non-coding DNA.

**Figure 1.**
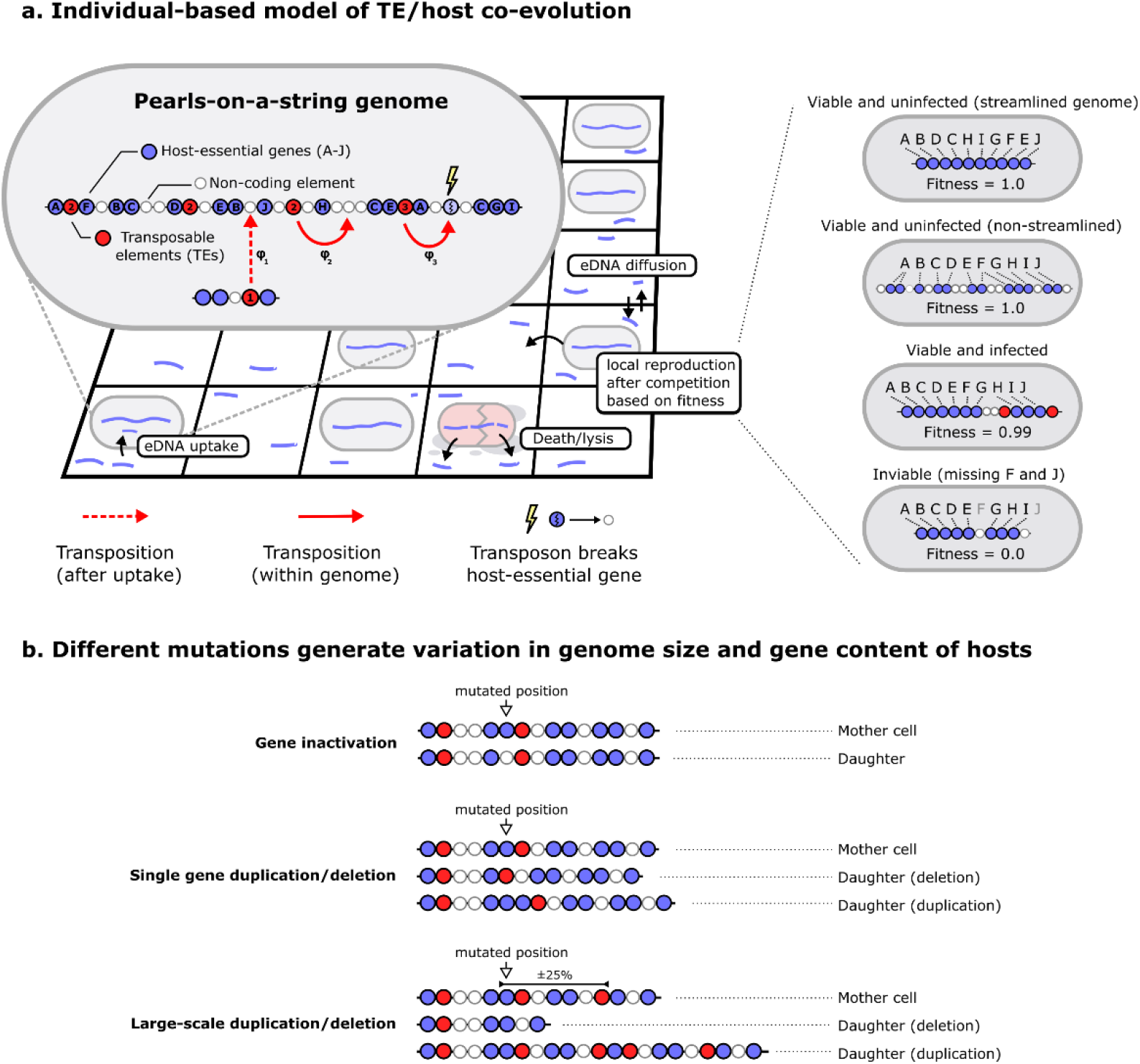
Individual-based model of co-evolving TEs and host genomes. **a.** Individuals are cells that undergo a process of birth, death, and DNA uptake on a spatial grid. Packaged within cells is a genome where genes are organised like “pearls-on-a-string” (Hogeweg et al. 2016). A total of 10 host-essential genes (A-J, in any given order) is necessary for cell viability. There is no explicit cost on the size of the genome, meaning that multiple (redundant) copies of genes may exist, as well as large stretches of non-coding DNA. The genomes also encode TEs that replicate through transposition independent of the host genome. Transposition of TEs happens both after uptake of eDNA (HGT) and during the lifetime of each cell. When transposons insert into to coding genes (a host-essential gene or another TE), that gene is inactivated and replaced by a non-coding element. To avoid TEs from growing indefinitely, a small fitness cost (c=0.005) is associated with each TE. The transposition rate of TEs is determined by φ, and may differ amongst individual TEs. **b.** Different mutations are depicted for cartoon genomes. Genomes are scanned from left to right upon reproduction, and each position may undergo mutation (illustrated in the cartoon with a white arrow). Mutations generate variation in genome size and gene content of individual cells. Large-scale duplications and deletions affect, on average, 25% of the genome and ensures genomes do not expand indefinitely (see Fischer et al. 2014). As mutations also operate on the level of TEs (i.e., modifying φ), the resulting model describes a multi-level coevolutionary process.

We assume that a maximally streamlined genome, *i.e.*, a genome encoding only one copy of each essential gene, has the same fitness as a genome that has multiple gene copies and large stretches of non-coding DNA. Fitness differences arise via differences in TE-abundance and site of TE insertion. Insertion of TEs into non-coding DNA, or redundant copies of host-essential genes is relatively harmless incurring a fitness cost of just 0.005 per TE. Insertion into essential genes (lightning symbols in **Figure 1a**) is lethal (fitness=0.0).

At each time step, cells compete locally for space. Space is a limiting resource that becomes available through cell death (see below). When a grid point is empty, competition occurs between up to eight cells from the neighbouring grid points with the winner being chosen at random, but weighted by fitness. The genome of the winning cell is replicated (with mutations) and the daughter cell is placed on the empty grid point.

TEs replicate independently of host genomes. If the rate of transposition (*φ*) is high, host-level selection struggles to prevent accumulation of TEs and ultimately hosts are driven extinct. When *φ* is low, TEs replicate too infrequently to compensate host-level selection and degradation from the eDNA pool. Ultimately, coexistence of TEs and hosts requires the possibility that TEs infect naïve (uninfected) lineages.

After the reproductive phase, non-viable cells plus a small fraction (*d*=0.02) of the healthy population, die. Dead cells lyse, spilling fragments of genome into the environment giving rise to a pool of extracellular DNA (eDNA) that can be taken up in a transformation-like process by the next generation of cells. Uptake happens at a fixed rate (*u*=0.01) and integration occurs with the same rate *φ* that determines transposition within genomes. Each time TEs replicate there is a small chance that this *φ*-parameter changes. Taken together, the model contains multiple levels (TEs within cells within spatially separated populations), with mutation and selection operating on each level.

### Spatial structure allows TEs and hosts to co-exist

Two important factors affecting the stable maintenance of cells and TEs are the degree of genome streamlining and rate of TE transposition. The manner in which these two traits interact depends on the scale of interactions, and particularly on whether or not interactions are confined to near-neighbours. To explore parameter space, mutation rates were first set to zero, and simulations performed over a range of fixed values of transposition rate (*φ*) and the degree of genome streamlining (ratio of host-essential to non-essential DNA).

As shown in **Figure 2a**, spatial structure – and thus local interactions – promotes coexistence of TEs and cells over a range of intermediate levels of genome streamlining and TE transposition rates (*φ*) (white points). TEs are unable to persist in cells that have highly streamlined genomes (high proportion of host-essential DNA) (blue inverted triangles), while cells containing less streamlined genomes are susceptible to extinction by TEs (red triangles).

**Figure 2.**
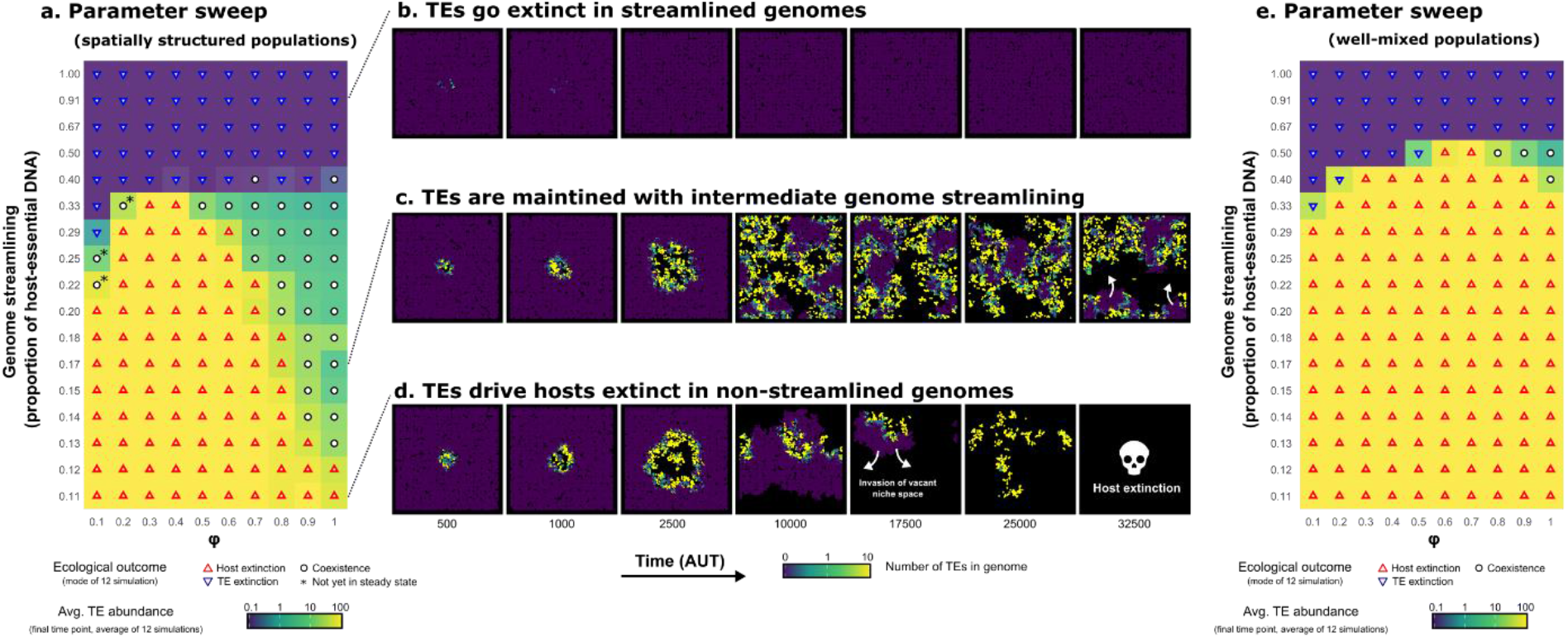
Spatial structure extends opportunity for coexistence of TEs and hosts at intermediate levels of genome streamlining. **a.** Heatmap shows the results of simulations that explore the relationship between the extent of genome streamlining, rate of transposition (φ), and coexistence of TEs and hosts. Simulations marked with an asterisk show apparent coexistence, which after 50,000-time steps have not reached a steady state (see the time courses of TE-abundance for both heatmaps in **Supplementary Figure 1**). Background colours indicate the average TE abundance in the final time points of the simulations, ranging from zero (purple) to 100 (yellow) on a log scale. **b-d**. Visualisations of TE abundance in spatially structured populations where b) TEs are rapidly lost, c) TEs stably coexist with their host, or d) TEs drive the population to extinction. Colours indicate the number of TEs in genomes, ranging from zero (purple) to one (green) to 10 (yellow). **e.** Heatmap similar to the one shown in a., but in well-mixed populations. For this, all individuals are assigned a random position on the grid after each round of replication.

The conditions for coexistence shown in **Figure 2a** depend on the interplay between three factors: i) genome streamlining, lethal mutations through TE insertion, and iii) the abundance of TEs in the (local) eDNA pool. In cells containing streamlined genomes, infection by TEs is likely to be lethal and thus there is little opportunity for the TEs to increase in frequency (**Figure 2b**). At the other extreme, cells with non-streamlined genomes are less likely to be killed by TE infection. This allows the TEs to increase in abundance, firstly within cells, and secondly in the eDNA pool. The latter then contributes to further amplification of TE abundance via HGT. As the TE load increases there is increasing chance that all cells become infected and from this state there is no possibility of recovery (**Figure 2d**).

Essential for the long-term survival of both TEs and cells, is the maintenance of uninfected cells (purple space in **Figure 2c**). This pool of uninfected cells decreases in frequency when infected by nearby TE-carrying strains, but can increase in frequency by recolonising vacant niche space made available by extinction events (white arrows in **Figure 2c**). Uninfected cells are then available for reinfection, resulting in a time-dependent cyclical process with chaotic waves (also see **Supplementary Video 1**). The critical factor for maintenance of both TEs and hosts is the time-to-extinction (and subsequent cell death and lysis) after infection. If this is too rapid (as happens with highly streamlined genomes), then TEs have little opportunity to amplify within genomes and eventually go extinct. If extinction is too slow, then all cells become infected and all host cells eventually go extinct.

Another important factor determining coexistence is the interaction range. If the infection of one strain readily infects another cell, the uninfected pool of cells is reduced over time. As evident from **Figure 2b-d**, spatial structure limits the spread of TEs to the local neighbourhood, which is likely important for co-existence. To test this, the simulations were repeated in well-mixed populations, where individuals are assigned a random position after each round of competition. The results show a highly significant reduction in conditions promoting coexistence (**Figure 2e**), thus demonstrating the central importance of spatial structure and local interactions.

### Genome streamlining evolves *de novo* in a structured environment

An intriguing finding from the above analysis is that streamlined genomes are resistant to invasion by TEs. This is evidently a lineage-level effect. To individual cells with streamlined genomes, infection by a TE is invariably lethal. In other words, what is costly to the individual appears beneficial at the lineage level. A central issue is whether this apparent example of altruism can evolve *de novo*.

To this end we introduced TEs into an evolving population of cells containing (initially) non-streamlined genomes. Mutations occur after each replication step, modifying the genome size and genome content of hosts, as well as the transposition rates of TEs (*φ*) (see Methods). Cells in the initial host population are all identical, carrying 10 host-essential genes and 30 non-coding positions, and were locally inoculated with TEs (in the middle of the grid).

Data in **Figure 3a** show that host genomes initially expand, but eventually evolved to be more streamlined. Three distinct episodes are notable. Initially (episode I) genomes expand in size. This is a consequence of TE amplification, but also entails an increase in the number of host-essential genes and non-coding elements. After expansion, a period of genome streamlining occurs (episode II). During this phase a decrease in the number of TEs and non-coding DNA is observed. The decrease in TE-abundance does not reflect a decrease in the transposition rate (*φ*), which instead increased over evolutionary time (**Figure 3a**, inset). Thus, TEs did not adapt to their host by becoming less infectious. Eventually, TEs and the amount of non-coding DNA reach a stable equilibrium (episode III), where genomes are comprised primarily of host-essential DNA (**Figure 3b**). Concurrent with genome streamlining is a decrease in vacant niche space (black areas in **Figure 3c**), which reflects decreased TE-driven extinction events and therewith an increase of the total population size (also see **Supplementary Figure 2**). In the steady state, most of the population consists of uninfected host cells, with occasional bursts caused by TE infection (**Figure 3c**, **Supplementary Video 1**).

**Figure 3.**
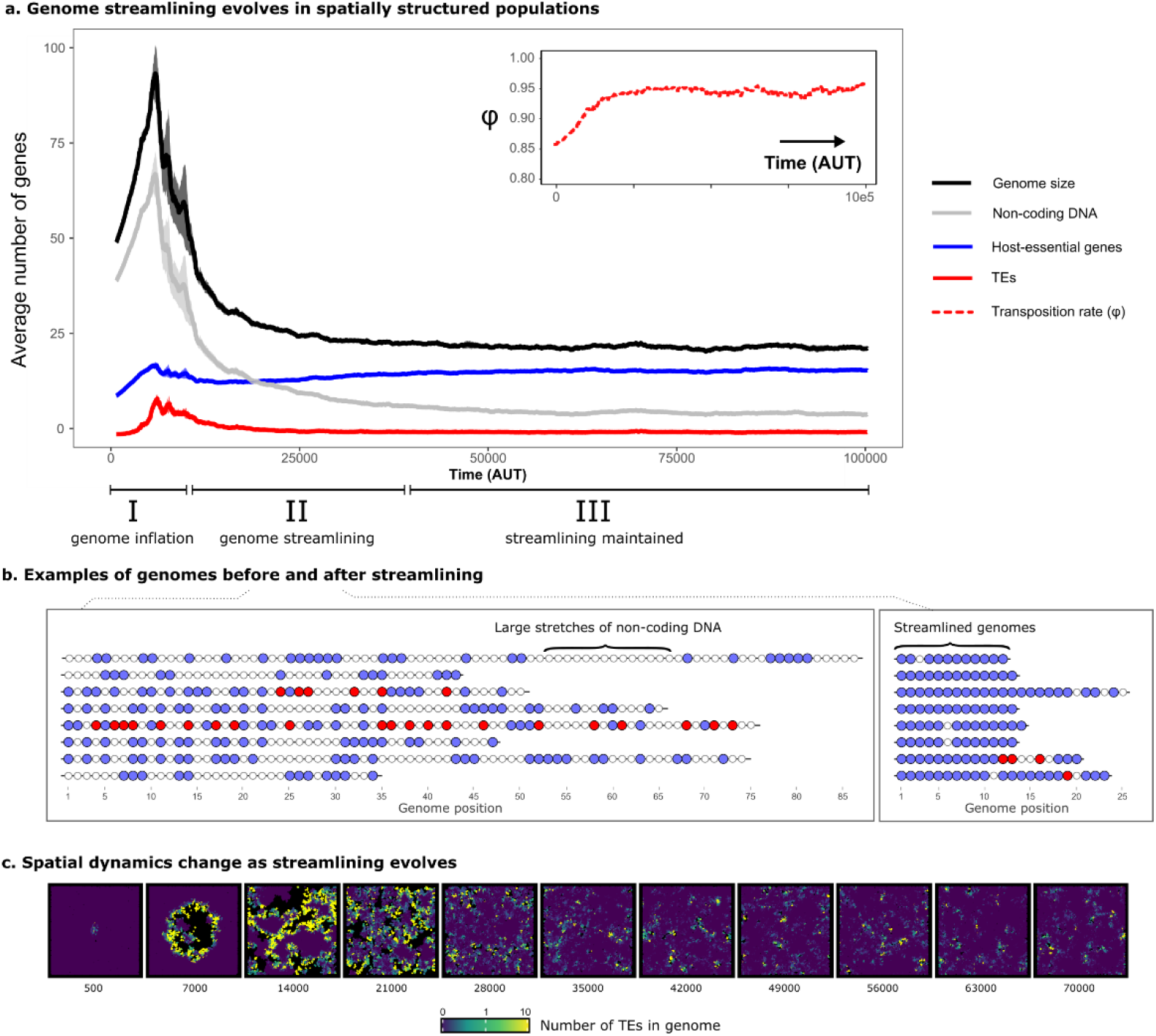
Co-evolution of TEs and host genomes drives genome streamlining. **a.** The average number of genes and genome size are plotted over time. Lines show the average of five independent simulations. Shaded areas denote the standard error across simulations. Host-essential genes are shown in blue, TEs are shown in red, and non-coding positions are shown in grey. Note that TEs persist (i.e. the red line is not zero) The inset shows the evolution of the average transposition rate (φ). The range of three distinct phases (I, II, and III) in the evolution towards streamlined genomes are shown along the x-axes. **b.** Examples of genomes in populations from panel **a**, before and after streamlining has evolved. Blue, red, and white dots denote host-essential DNA, TEs, and non-coding DNA respectively. **c.** TE-abundance per individual, which correspond to equally spaced time points from a single population from panel **a**. A log-transformed gradient of blue to green to yellow indicates increasing numbers of TEs inside individual genomes. Black indicates empty space where cells have locally died out, which provides niche space for invasion by other lineages.

As evident in the ecological simulations above (see **Figure 2**) the time taken for lineages to go extinct is an important factor and is expected to evolve during the course of the selection experiment. The mechanistic nature of the model means that individual cells can be retrieved after the simulation completes, and their evolutionary history directly observed. **Figure 4a** shows, for each extinct lineage, the number of TEs that accumulate from the time of TE infection until extinction. Lineages of cells with non-streamlined genomes (before streamlining evolved, generation 100 to 400) persist for longer after infection and liberate many more TEs into the eDNA pool compared to lineages of cells with streamlined genomes (generation 1,800 to 2,100). The histograms from **Figure 4b** show that streamlined genomes go extinct rapidly (no more than a few generations), and as a consequence only produce a few TEs (**Figure 4c**). Thus, although streamlined genomes produce fewer progeny in the short term, they eventually shape an environment in which they thrive. Moreover, after non-streamlined genomes have succumbed to bursts of TE infection, streamlined genomes are free to invade the space freed by cell lysis (**Figure 4d**).

**Figure 4.**
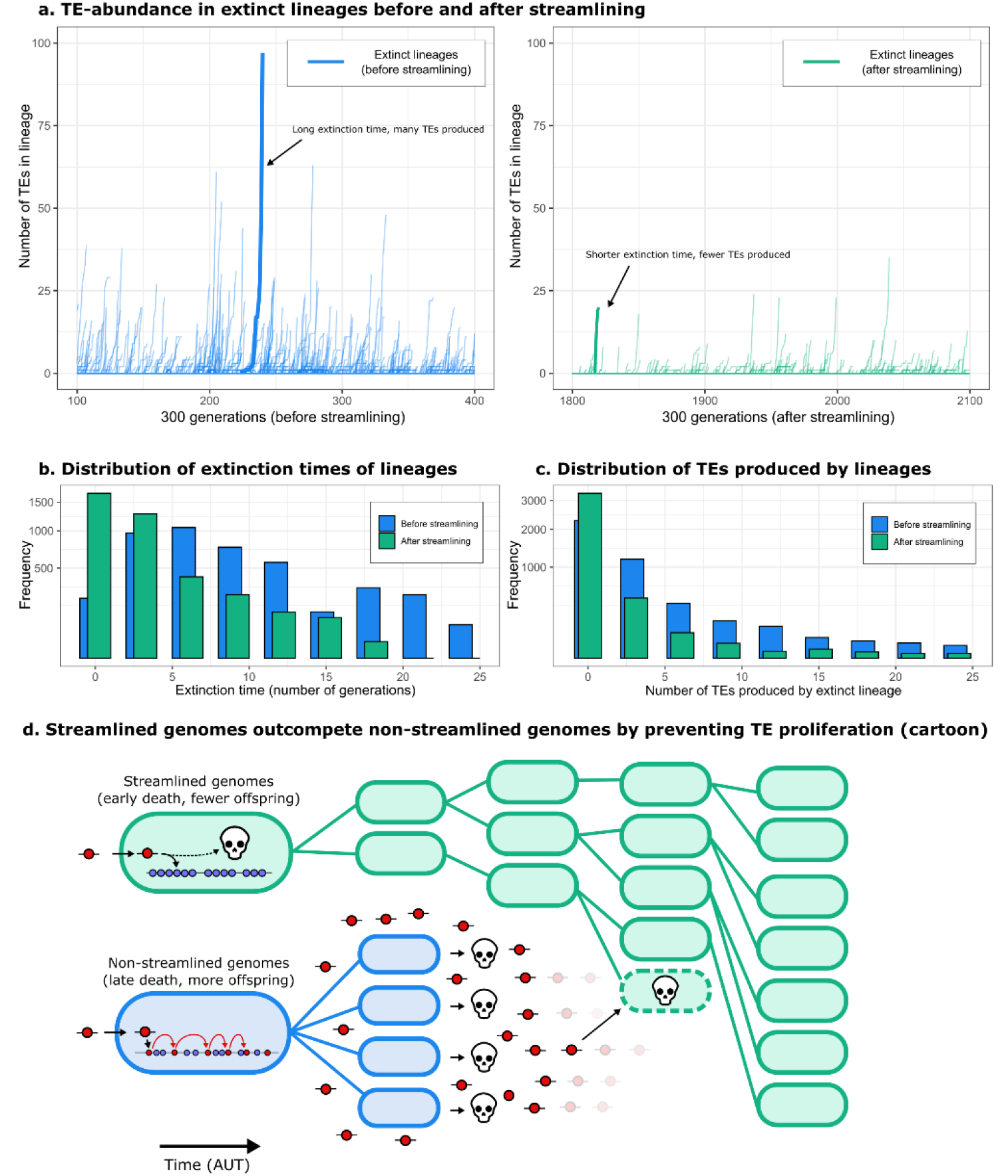
Genome streamlining reduces extinction time and hampers TE proliferation. **a.** Before genome streamlining, infected lineages produced more TEs until they went extinct. The x-axis shows the generation number, and the y-axis the number of TEs (for each extinct lineage during this time interval). The left-hand side shows lineages of cells before streamlining evolved (in blue), and the right-hand side after streamlining has evolved (in green). Two arbitrary lineages are highlighted with a thick line for illustrative purposes. **b-c.** Histograms for the extinction time (number of generations since infection) and the number of TEs produced by extinct lineages, before and after streamlining (blue and green respectively). Note that the y-axes are square-root transformed to clearly illustrate the difference between the two distributions. **d.** Cartoon illustrating how cells containing streamlined genomes (green), despite spawning fewer progeny in the short term when infected by the same number of TEs, eventually replace cells containing non-streamlined genomes (blue) by limiting opportunities for TE proliferation. Although streamlined genomes may be infected by TEs derived from non-streamlined genomes, newly acquired TEs have little opportunity to amplify because infection of a cell with a streamlined genome is invariably lethal.

### Local interactions are essential for the evolution of genome streamlining

The data shown in **Figure 3c** indicates an important role for spatial structure. To test this directly, we repeated the simulations, but assigned individuals to a random position after each round of competition and reproduction. Starting from conditions that are ecologically viable according to **Figure 2** (10 host-essential genes and 10 non-coding elements), populations rapidly evolved larger genomes but were eventually driven to extinction by TEs (**Supplementary Figure 3a**). Similar results were obtained when mixing was confined to just the eDNA pool (**Supplementary Figure 2b**) or when TEs were not amplified by within-cell replication, but exogenously delivered (**Supplementary Figure 3c**). Moreover, populations that had already evolved genome streamlining (from previous experiments) rapidly went extinct when the local interaction neighbourhood was removed by mixing (**Supplementary Figure 3**). Local interactions and direct feedback from the environment are thus essential for the evolution and maintenance of streamlined genomes.

### Genome streamlining is driven by TE-induced inactivation of host-essential genes

The evolution of genome streamlining appears adaptive in that it results in the elimination of TEs – at least temporarily – from local populations. But to individual cells, genome streamlining is clearly maladaptive: infection of cells with streamlined genomes is invariably lethal. The evolution of genome streamlining thus appears to be attributable to selection at the level of lineage viability, with those lineages comprised of streamlined genomes outcompeting lineages with less streamlined genomes (as illustrated in **Figure 4d**).

To test the hypothesis that lineages of cells containing streamlined genomes gain a lineage-level benefit that derives directly from the cost experienced by individual cells, we repeated the above simulations but denied the possibility that TE movement causes lethal effects. Under these conditions, genome streamlining no longer evolved (**Supplementary Figure 4**). TEs and their hosts nonetheless persist through continuous waves of infection and recolonization of available niche space. This result demonstrates that the evolution of genome streamlining (**Figure 3**) is driven by TE-generated mutations that are harmful to individual cells.

### Persistence of TEs depends on rock-paper-scissor dynamics

Given that cells containing streamlined genomes drive TEs extinct, the persistence of TEs shown in **Figure 3c** seems counter-intuitive. However, understanding emerges from examination of the eco-evolutionary dynamics (see **Supplementary Video 1**), combined with observation of the evolution of streamlined genomes in the absence of TEs. Starting with the latter, data in **Supplementary Figure 5** shows that non-streamlined genomes replace streamlined genomes in the absence of TEs. This occurs in part as a consequence of duplication bias (in a minimal genome, deletion-mutants are never viable – making duplications the only mutations that change genome size), but also because genome expansion generates multiple copies of essential genes that confer mutational robustness. Thus, in the absence of TEs, selection favours larger genomes.

In data from evolutionary simulations (**Figure 3**), TEs are never absent. Instead, they decline to low numbers in local patches and once rare, individual cells with non-streamlined genomes are favoured over cells with streamlined genomes. Apart from the selective benefits of larger genomes described above, this also occurs because non-streamlined genomes are, at least initially, less sensitive to the deleterious effects of TE infection. However, as the load of TEs within lineages increases, costs are increasingly realised at the level of local lineages. This then establishes conditions that once again favour the evolution of cells with streamlined genomes. Cells with streamlined genomes thus beat TEs, which beat non-streamlined genomes, which beat streamlined genomes, and so on, in a cyclical game of rock-paper-scissors (**Figure 5**). The long-term persistence of TEs and cells with streamlined genomes depends on this dynamic. As the re-emergence of non-streamlined genomes entails evolution, disabling mutation in populations which evolved streamlined genomes breaks the rock-paper-scissors cycle, eventually driving TEs extinct (**Supplementary Figure 6**).

**Figure 5.**
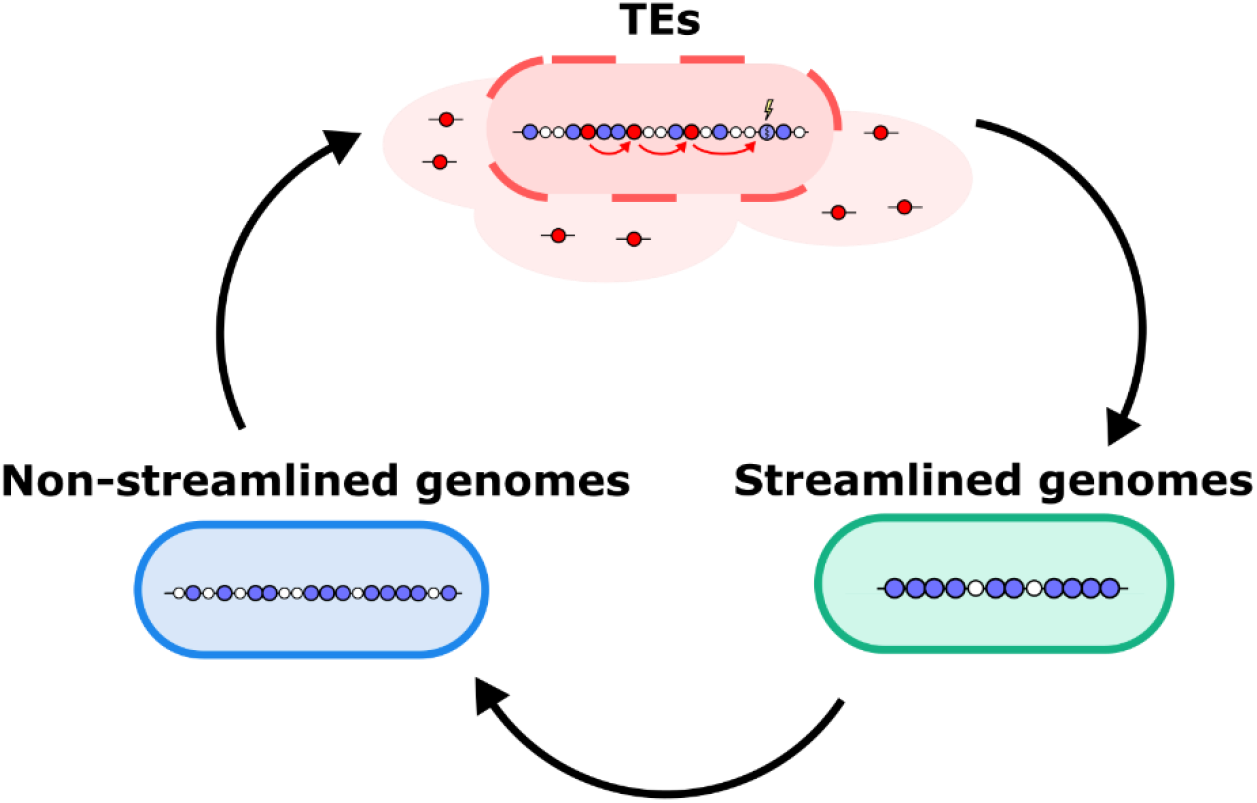
Rock-paper-scissors dynamics allows TEs and cells with streamlined genomes to coexist. A cartoon illustrating how both streamlined genomes and TEs can be maintained within the population indefinitely. As streamlining lowers the (local) abundance of TEs, non-streamlined genomes are favoured. This enables TEs to once again infect cells and locally thrive, which in turn upholds the selection pressure for streamlined genomes.

### TEs do not drive genome streamlining in sexually reproducing populations

As TEs require transfer to new linages to persist, simulations in which DNA uptake is disabled result in TE extinction (**Supplementary Figure 5**). However, TEs in nature can also persist in populations through sex and recombination. Our model was therefore modified to incorporate sexual reproduction by disabling DNA uptake and implementing a simplified form of sexual reproduction. In these populations, competition for vacant space is determined by sampling two individuals from the local neighbourhood (weighed by their fitness), and their genomes recombined via a single cross-over event (see Methods). Importantly, this process allows TEs to infect new lineages without transposition, removing the risk of lethal mutations. These sexually reproducing “eukaryotic” populations (**Figure 6** purple lines) did not evolve genome streamlining and grew large in size compared to prokaryotic populations (**Figure 6**, green lines). Accordingly, the average fitness of sexual populations is relatively low, as a substantial fraction of the population was infected with a large number of TEs. In the absence of sex and horizontal gene transfer (*i.e*., in strictly clonal populations) TEs went extinct and genome size increased (**Figure 6**, blue lines).

**Figure 6.**
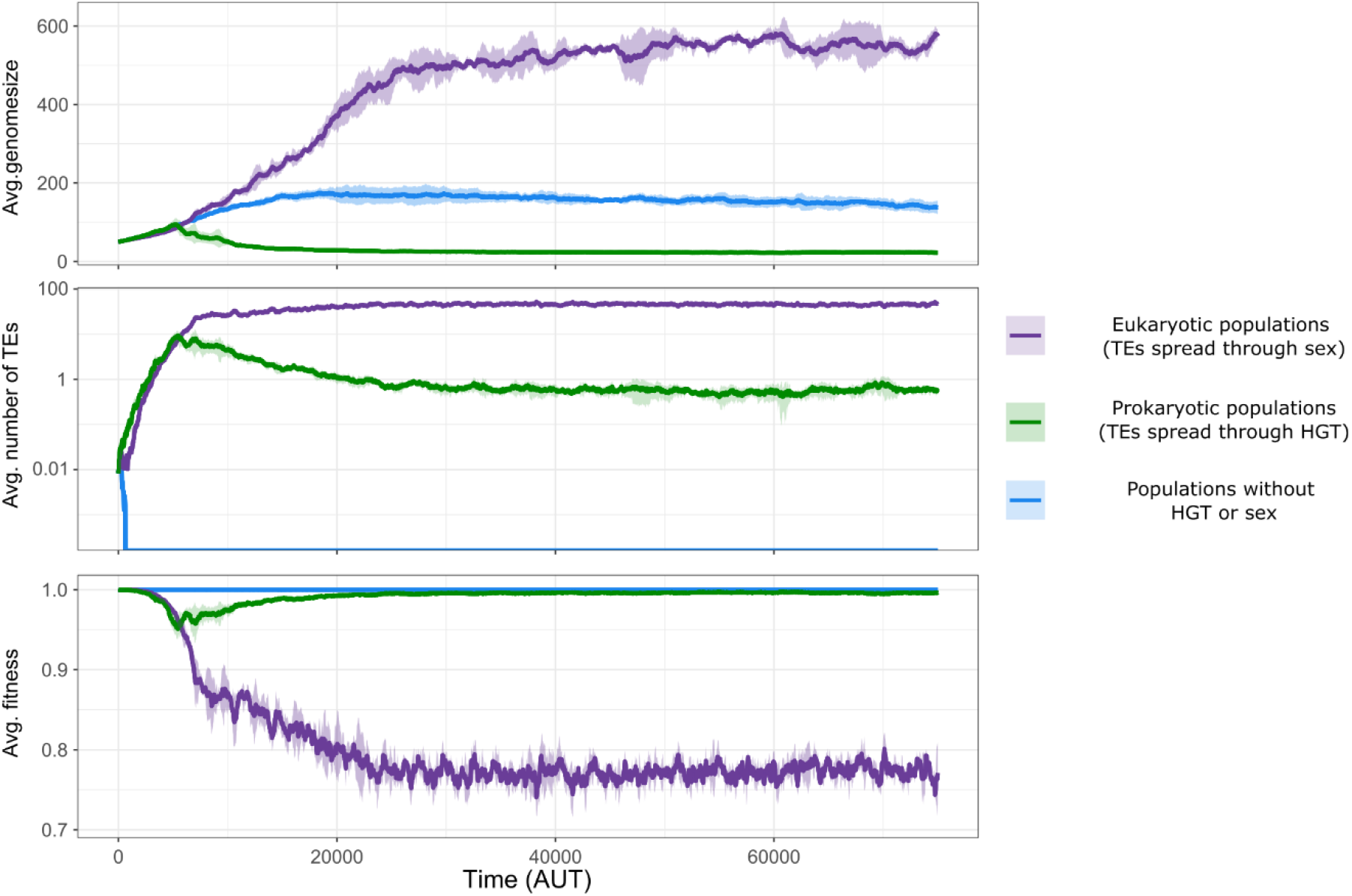
Streamlining does not evolve in sexually reproducing populations because recombination partially unlinks TEs from the deleterious effects they cause. For three different population types, the genome size, TE-abundance, and average fitness are shown. Shaded areas are the standard errors from multiple independent simulations. Prokaryotic populations are shown in green (n=5), sexual populations in purple (n=3), and populations without HGT or sex in blue (n=5).

To understand the absence of genome streamlining in sexual populations, it is necessary to reconsider the two steps of successful TE amplification. First a TE must infect a host, but in order to increase in frequency, it must also replicate within the host (at least once). In the asexual populations, cells with streamlined genomes die during the infection step, thus immediately blocking further TE amplification. With recombination, however, TEs can infect new lineages without risk of immediately killing the host, irrespective of the level of genome streamlining. Although subsequent transposition may still render the host inviable, the host cannot prevent a TE from infecting its genome, thus allowing the TE to replicate at least once. The fact that genome streamlining does not occur in these sexual populations suggests that streamlining in asexual populations evolves to prevent transposition *between* genomes, and not transposition *within* genomes.

## Discussion

Here we have presented an *in silico* co-evolutionary model of transposable elements (TEs) and host genomes. The model reveals an interesting interplay between genome streamlining (the amount of coding DNA) and TE-abundance. Selection initially favours cells with expanded genomes, because additional genome space reduces the chance that transposition has deleterious effects. However, while adaptive at the level of individual cells, cells with expanded genomes provide opportunity for the population of TEs to increase in the lineage of descendent cells to the point where extinction of the lineage becomes inevitable. When the environment is spatially structured, extinction is localised, and the resulting vacant niche-space creates opportunity for cells with streamlined genomes to invade. As TEs cannot replicate within streamlined genomes without killing their host, this leads to extinction of the local pool of TEs. Absence of TEs eliminates the selective advantage for streamlining and thus genomes expand in size. These interactions cycle generating rock-paper-scissors dynamics, where TEs beat non-streamlined genomes, streamlined genomes beat TEs, and non-streamlined genomes beat streamlined genomes. In sexually reproducing populations, streamlined genomes have no advantage over non-streamlined genomes because recombination unlinks TE-infection from potential DNA damage. Thus, our co-evolutionary model of TEs and hosts provides an explanation for streamlined genomes in prokaryotes, and expanded genomes in eukaryotes.

Interestingly, genome streamlining is maladaptive at the individual cell level, but is selectively favoured because of benefits that accrue to lineages of cells. This behaviour, which is costly to individual cells, is analogous to abortive infection, a well-studied mechanism that protects cellular collectives against bacteriophages (Lopatina et al. 2020). Earlier modelling on spatially structured populations has already illustrated that early death can be favoured when it promotes the long-term survival of the lineage (Boerlijst et al. 1993; Werfel et al. 2015). Our results connect these observations to the evolution of genome architecture, showing that early death is an evolutionarily attainable (and maintainable) protection mechanism against TEs.

For illustrative purposes, we have deliberately not included other mechanisms that are known to result in genome streamlining. For example, alternative hypotheses for the different structures of prokaryotic and eukaryotic genomes are differential energy budgets (Lane & Martin 2010), deletion biases in prokaryotes (Kuo & Ochman 2009; Sela et al. 2016), and the small population sizes of eukaryotes (Lynch 2006). For prokaryotes in particular, recent studies have suggested that natural transformation may play an important role in the removal (rather than the acquisition) of mobile elements (Croucher et al. 2016; Apagyi et al. 2018). Although we show that our mechanisms can operate in isolation, it is likely that these interplay with a range of additional factors.

The TEs in our model are based on insertion sequence (IS) elements, a particular yet common class of TEs. TEs are assumed to be autonomous (they encode their own transposase function), show no notable bias in insertion site preference, and move both vertically and horizontally. Not all TEs are marked by these characteristics. For example, REPIN sequences (**r**epetitive **e**xtragenic **p**alindromic sequences forming a hairp**in**) take up many intergenic spaces in *E. coli* and *Pseudomonas fluorescens* SBW25 (Bertels & Rainey 2011; Park et al. 2021). However, REPIN sequences do not move autonomously and are replicated by a single-copy transposase that has been vertically inherited for millions of years (Bertels & Rainey 2011). The pattern of REPIN sequence abundance may therefore be explained by a direct fitness advantage that would evidently not promote genome streamlining.

Questions unanswered through use of *in silico* models concern relevance to the natural world. However, there are reasons to assume the likelihood of legitimate connections, particularly given the spatially structured nature of microbial populations, the abundance of TEs, and pervasiveness of HGT. Evidence could be sought by interrogation of genome sequences from a set of phylogenetically related strains sampled at precise spatial and temporal scales. However, the dimensions of those scales would firstly require experimental investigation. An alternate possibility is to compare data on the relationship between genome size and TE-abundance derived from the analysis of diverse genome sequences with theoretical predictions of this relationship at equilibrium. The latter can be derived from our model populations by analysis of the genomes of all viable cells present at the end of the simulations. The data, shown in **Supplementary Figure 7**, indicate a strong positive correlation between genome size and TE-abundance only under conditions where TEs cause harmful effects. Precisely such a relationship has been previously reported (Touchon and Rocha 2007) for IS-elements, with the authors even suggesting that such a relationship might reflect robustness of larger genomes to lethal TE insertion.

In general, our model is not sensitive to the precise choice of parameters. We consistently observed the streamlining of genomes and long-term stability when i) TEs and their host coexist, ii) the rate of transposition-induced inactivation is greater than zero, and iii), the spatial grid is large enough to allow major bottle necks in population size. This long-term stability can be explained by rock-paper-scissors dynamics (Rainey & Travisano 1998; Kerr et al. 2002; Ferguson et al. 2013), similar to what is observed for susceptible-infectious-recovered (SIR) models (van Ballegooijen & Boerlijst 2004). Besides choices in parameters, there are however also important assumptions our model makes that warrant future exploration. For example, it would be interesting to extend the model to include genes that are not essential but still contribute to the fitness of the organisms. Earlier work has shown that the uptake of non-essential genes is adaptive to cells, even in the presence of selfish genetic elements (van Dijk et al. 2020). However, the accumulation of non-essential genes likely trades off against the protection against TEs as illustrated in this study. As trade-offs are a well-known mechanism for diversification, these are exciting angles for future work.

## Author contributions

BD conceived the model with input from NT. BD designed and/or programmed the model, ran simulations, generated figures from the output data, and drafted the initial manuscript. FB, NT, LS and PR contributed ideas and concepts that shaped the design of in silico experiments and crafting of the manuscript. All authors gave final approval for publication and agree to be held accountable for the work performed therein.

## Acknowledgements

I (BD) thank Paulien Hogeweg, Jeroen Meijer and Hilje Doekes for fruitful discussions during the early stages of the project.

## Methods

The model implemented in this study is an individual-based model (IBM) of the co-evolution of transposable elements (TEs) and their host. It is composed of a spatial grid with i) simple cells that compete and reproduce; ii) naked DNA from prior generations which is taken up by cells (see **Figure 1** in main text). Cells carry simple genomes with three distinct genetic elements: host-essential genes, transposable elements (TEs), and non-coding DNA (see section: Genome structure and fitness). TEs are modelled as selfish genetic elements which replicate within genomes, and integrate after DNA uptake, with rate *φ* (see section: TE-dynamics and HGT), and can infect new lineages through a transformation-like process. The gene content and genome size of the host cells can vary through mutations (see paragraph: Mutational processes) which occurs after competition and reproduction (see paragraph: Competition, and reproduction). As a consequence of mutation, each TE can also have a unique transposition propensity (*φ*). In short, the model contains multiple levels, describing both the ecology and the evolution of TEs and their host genomes.

### Genome structure and fitness

Individuals carry a genome that encodes a linear sequence of genetic elements. We assume that cells need to perform 10 essential functions, and therewith there exists *k* essential genes (A-J). We assume that these essential functions are performed when an essential gene with function *i* is present at least one (e_i_> 0). Carrying multiple copies of these genes does not directly impact fitness (*f_i_*). However, a genome that lacks one of these essential functions has fitness zero, meaning it cannot (or no longer) compete for reproduction and dies in the next time step. The second type of genetic elements we consider are TEs, which self-replicate within genomes. The total number of TEs (T) confer a small cost (c) to the host. The fitness of the host then becomes:

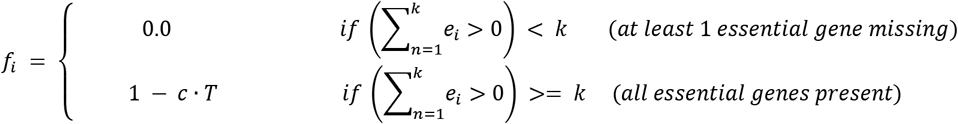

Note how the third class of genetic elements, non-coding DNA, does not impact fitness. Although it can be generated, amplified, or trimmed through mutational processes, we deliberately avoid implementing costs for genome size to illustrate how TEs drive the streamlining of genomes.

### Mutational processes

Mutations happen every time genetic elements are replicated, and change the gene content and genome size of individuals. When cells reproduce, their genomes are scanned from left to right, allowing each genetic element to undergo mutations once. Changes can be applied to the genetic element itself, or they can be the start site of a large-scale deletion, duplication, or inversion of multiple genes. These large-scale events enable the rearrangement of gene order, and also ensure that genomes do not grow indefinitely in the absence of a deletion bias (Fischer et al. 2014). Single genetic elements can be deleted, duplicated, or inactivated (transforming host-essential genes and TEs into non-coding genes). TEs can also change their transposition propensity *φ* with a uniform step size (0.1) up or down.

### Death and lysis

Every time step, a small subset of the host population stochastically dies with probability *d*. Moreover, cells that are not viable (or no longer viable due to transposon-induced mutations) also die. Before being removed from the grid, dead cells spill their DNA into the environment. This DNA is uniformly fragmented into pieces of 3 to 8 genetic elements each. These stretches of DNA are taken up by future generations with a fixed rate *u*.

### TE-dynamics and HGT

The dynamics of TEs occurs through two distinct processes. Firstly, TEs can replicate during the lifetime of a cell with rate *φ.* Note that one genome can carry multiple TEs, potentially with different *φ*-parameters. The second process by which TEs spread is by means of a transformation-like process. Living cells take up naked DNA derived from prior generations, after which TEs can integrate into the host chromosome with the same rate *φ*. We assume that both transposition events (i.e. after uptake or during the cell’s lifetime) occur at random positions in the chromosome, and can cause the inactivation of genes at the insertion site (lightning symbols in Figure2a). When a gene at the site of transposon-insertion codes for one of the host-essential functions, this event is lethal for cells (unless another copy of that gene is still active). Transposons inserting into non-coding DNA never have a (direct) damaging effect.

In the main text, we also test what happens when inactivation of genes at the insertion site was removed. For these populations, TEs are always inserted “next to” rather than “into” the genetic element at the insertion-site. Note that whether TEs insert “into” or “next to” genetic elements at the insertion site does not influence their transposition dynamics, but only the potential damage that transposition may cause.

Our model assumes a TE can transpose directly after the uptake of DNA. Although transposition events from naked DNA have been shown to occur (Domingues et al. 2012) by using the machinery encoded on the TE itself (Kloos et al. 2020), our model does not explicitly assume this to be the only mechanism. An alternative route of HGT of a TE would be *via* an intermediate mobile genetic element such as a plasmid or phage (Mark Osborn & Böltner 2002; Pfeifer et al. 2021). The subsequent transposition to the host chromosome would still carry the risk of transposon-induced mutations. In principle, our model abstracts away from these distinctions.

### Competition and reproduction

Each time step, competition happens for unoccupied grid points. Up to eight cells in the direct (Moore) neighbourhood compete proportional to their fitness. The relative chance individual *i* wins this competition (*R*_i_) is determined by the individual’s fitness (*f_i_*), divided by the total fitness of all competitors (*f_j_*) plus a constant ∊. The latter constant ensures that a single individual does not win by default, and ensures that it is unlikely for any individual to reproduce when all competitors are unfit.

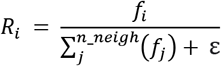

### Sexual reproduction

To distinguish the process of HGT from sex and recombination, we implemented a simple mode of sexual reproduction in our model. For this, two competitors are sampled proportional to their fitness (see above), and their genomes are recombined by a single cross-over event in the middle of their genomes. The resulting (haploid) genome undergoes mutations in the same way as a clonally reproduced genome would in the base model. These sexually reproducing populations do not take up environmental DNA.

### Parameter choice

Because our model is an abstraction of biological processes, it is not trivial to estimate the precise values that would be realistic/accurate. We however found that, given parameters that allow for host/TE coevolution, our main results are robust to the precise values of parameters. Apart from the parameter sweeps presented throughout this study, we therefore chose to parameterise the model by finding (biologically reasonable) parameter values where:

- **TEs can (potentially) coexist with their host genome** Requires sufficient HGT (or sex) for TEs to jump to new lineages Requires local extinctions such that healthy lineages can invade empty space (through a direct fitness-cost on the TEs, through lethal TE-insertions, or both) DNA diffusion is low, so that TEs can only infect local strains (and not the entire population at once)
- **The model remains computationally feasible** Large-scale duplications and deletions are assumed, so that genomes do not grow indefinitely (Fischer et al. 2014) The system size is set to the minimal size where local extinctions, wave fronts, and multiple strains can occur simultaneously

Many other variables (gene content, genome size, transposition rate) are allowed to evolve. See **Table 1** for a full list of parameters and their values.

**Table 1:** List of model parameters and values (unless stated otherwise)

| Parameter | Value | Description |
| --- | --- | --- |
| Grid size (W,H) | 150,150 | Size of (toroidal) grid on which individuals reside |
| Number of host-essential functions (k) | 10 | The number of unique functions that a cell must perform to be viable. (i.e. the minimal genome size is equal to k) |
| Fitness penalty per TE (c) | 0.005 | Per-TE penalty on fitness |
| Natural death rate (d) | 0.02 | Probability of stochastic death for individual cells |
| DNA uptake (u) | 0.01 | Probability of uptake per (local) fragment of DNA (after uptake, the chance of successful integration is determined by the ϕ-parameter of the TE) |
| Rate of transposition events within genomes (j) | 0.01 | Probability of transposition-event during the lifetime of a cell (after this even has been invoked, the chance of successful transposition is further determined by the ϕ-parameter of the TE) |
| DNA diffusion (D) | 0.01 | Probability of fragments to move into a random neighbouring grid point |
| TE transposition rate ($\phi$) | 0.9 / evolvable | Individual success-rate of transposition (can be unique/evolvable per TE) |
| Single-gene duplication/deletion | 0.001 per position | Per gene per generation probability of duplication or deletion |
| **Large scale duplication/deletion/inversion** | 0.001 per position | Per gene per generation probability of position invoking a large-scale duplication, deletion, or inversion |
| Gene inactivation | 0.001 per position | Per gene per generation probability of gene inactivation (position becomes non-coding) |
| TE transposition mutation | 0.01 per TE | For every TE replication (transposition, host-genome replication), the probability that the transposition rate changes with a uniform step size of $\pm 0.1$ |
| TE-induced damage | TRUE | When a TE inserts into the list of DNA elements ("the genome"), the adjacent coding genes get inactivated (converted into non-coding DNA) |

### Software used

The individual-based model presented in this study is a C++ extension of Cash (Cellular Automaton simulated hardware), originally written in C R.J. de Boer and A.D. Staritsk. All analyses were done in R, using the packages ggplot2 (Wickham 2016), dplyr (Wickham et al. 2015).

### Code availability

The code for the individual-based model and the scripts to run parameter sweeps are available on Github. (https://github.com/bramvandijk88/selfishDNA).

## Supplementary material

### Supplementary Figures

**Supplementary Figure 1.**
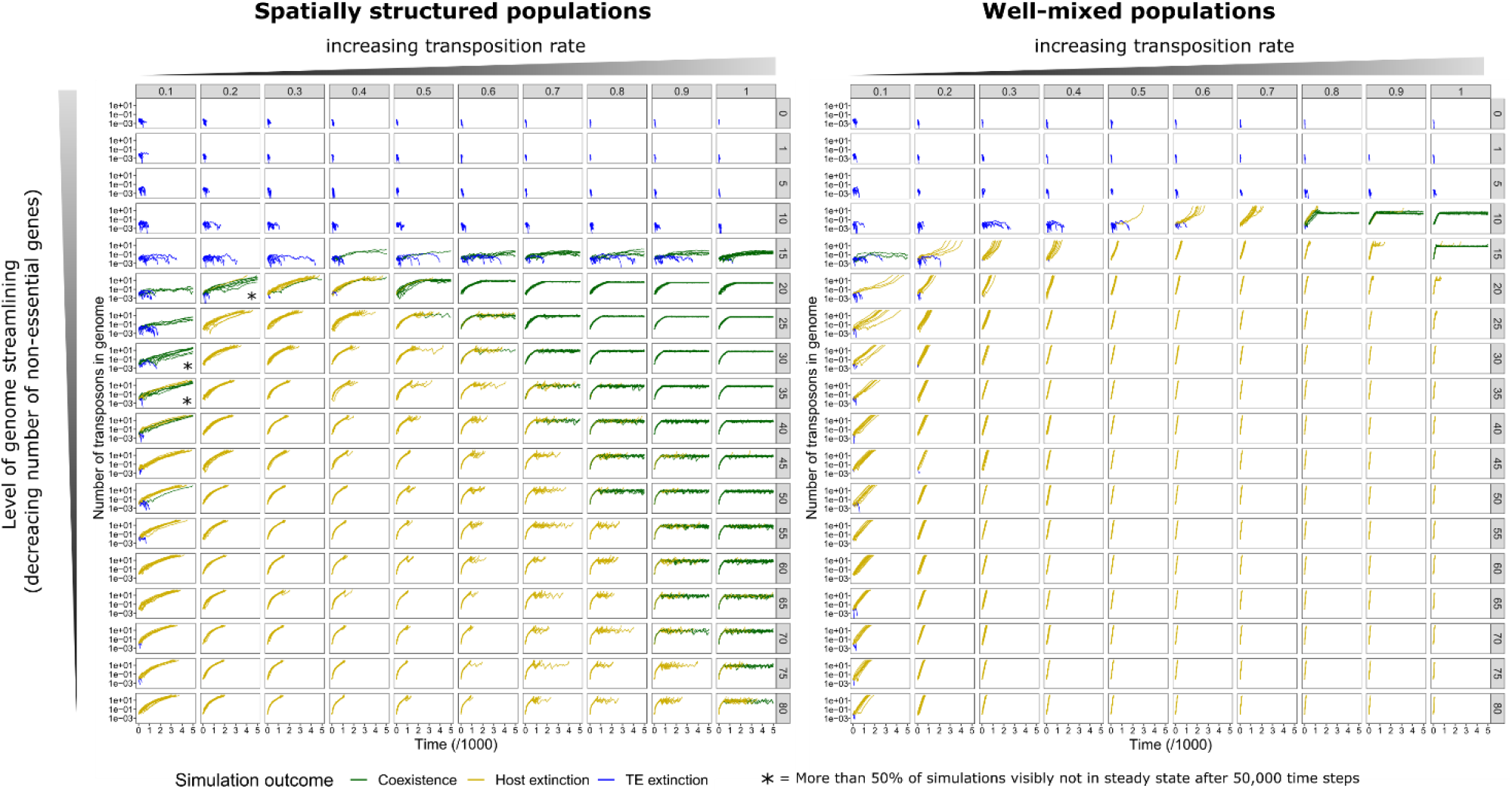
TE-abundance from parameter sweep in Figure 3 from main text. These are the raw data for the heatmaps shown in Figure 3a. 12 simulations were run for each combination of transposition rate (columns) and genome streamlining (rows). Each window shows the average number of transposons in genomes (y-axis) in the population over time (x-axis). The line colours denote whether the TEs went extinct (blue), the population went extinct (yellow), or the TEs coexist with their host until the end of the simulation (50,000 time steps, green). The data show that the spatially structured populations allow for stable TE abundances in steady state for many different parameters. At low transposition rates, these populations are clearly not yet in steady state, but appear to be going towards extinction. Coexistence is also possible in the well-mixed model, but only when the number of coding and non-coding genes is exactly the same (Park et al. 2021).

**Supplementary Figure 2.**
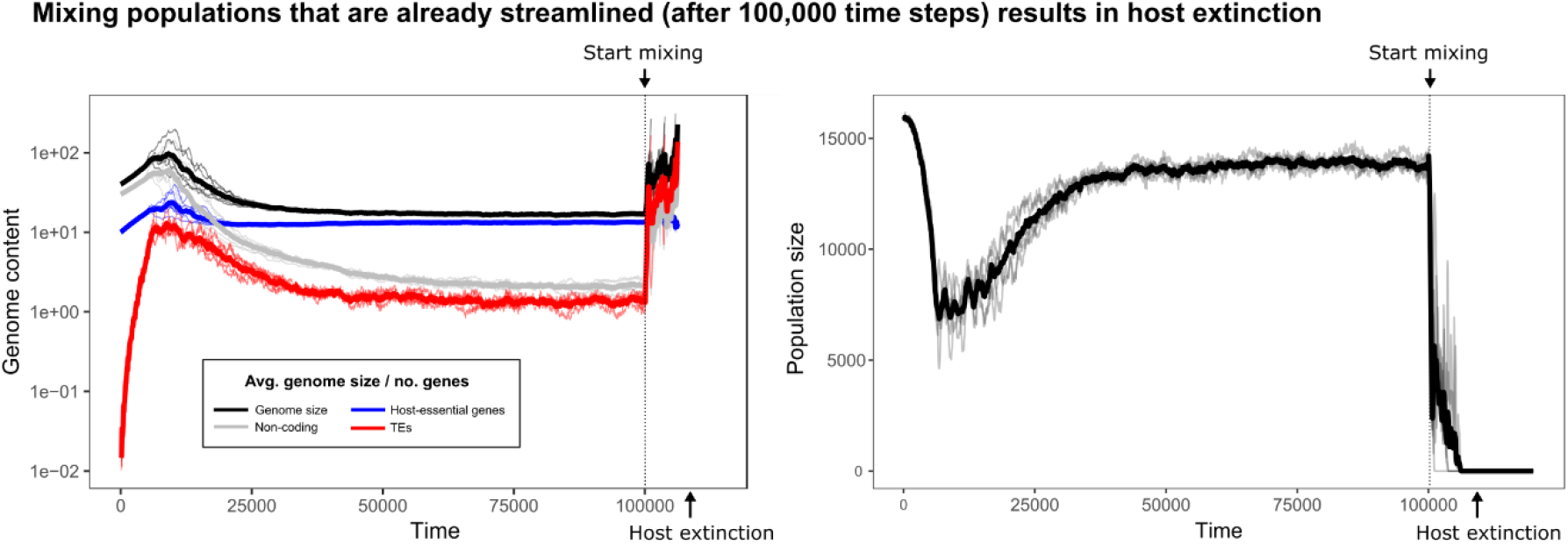
Mixing already streamlined populations (from T=100,000) results in extinction of the population. For five replicate populations where streamlining has occurred, the population was mixed continuously after 100,000 time steps. Despite the streamlined genomes, TEs are able to expand and drive the eventual extinction of the population. Thin lines denote the average genome size, gene counts, and the population sizes of individual simulations, whereas thick lines are the average across simulations.

**Supplementary Figure 3.**
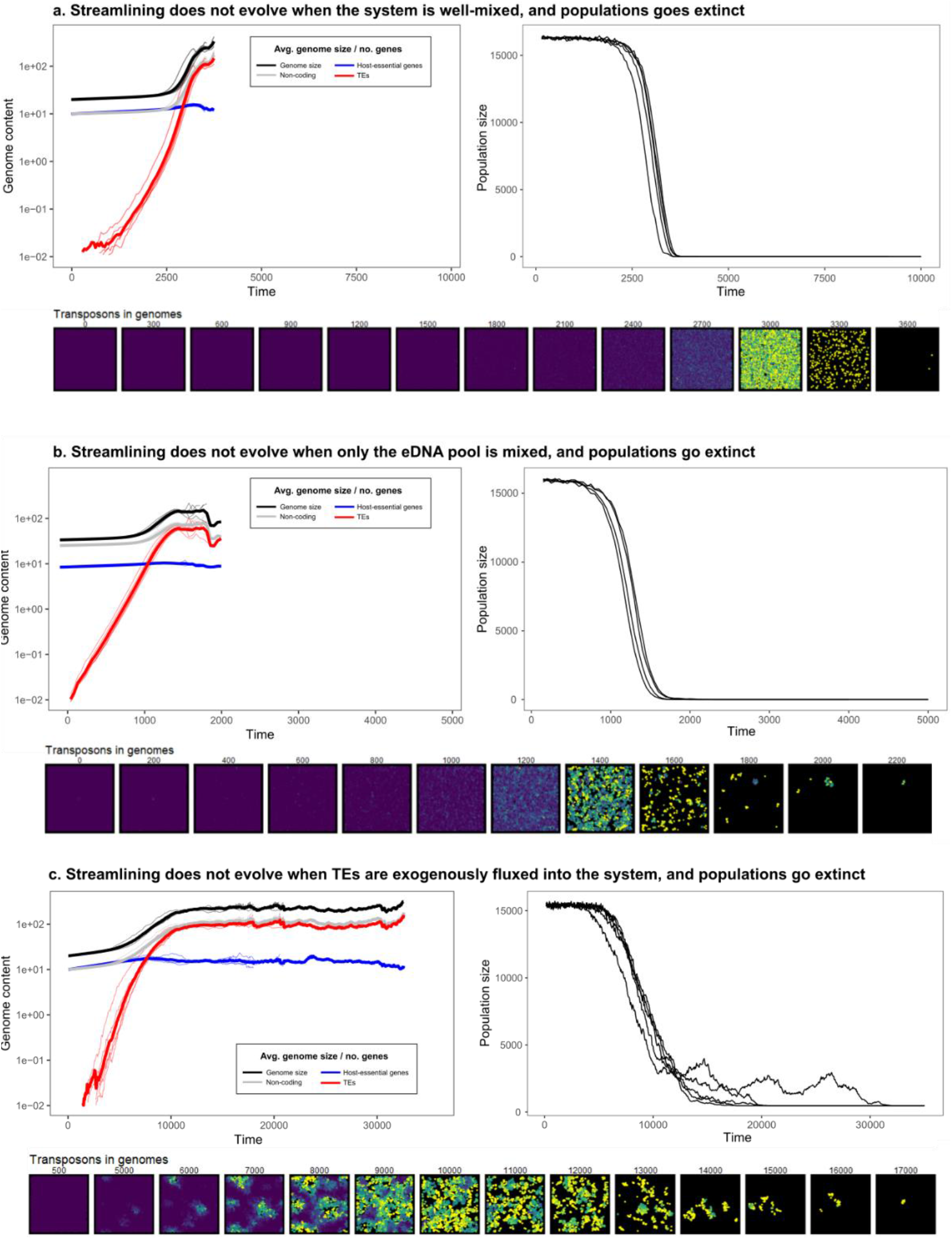
Streamlining does not evolve in well-mixed systems, when only the DNA pool is mixed, or then TEs are exogenously fluxed into the system. **a.** Starting from a population that ecologically supports coexistence (10 host-essential genes, 10 non-coding genes), well-mixed populations evolve larger genomes, and quickly go extinct. model with mutations quickly drives the evolution of larger genomes. **b.** Starting from a population that ecologically supports coexistence without mixing (10 host-essential genes, 30 non-coding genes), mixing the eDNA pool also results in extinction of the population. **c.** Starting from a population that ecologically resists TEs (10 host-essential genes, 10 non-coding genes), fluxing in TEs exogenously (instead of spilling them into the environment from dead cells), results in short-sighted evolution of large genomes. These large genomes survive longer when infected by TEs, but eventually go extinct. For subfigures a-c, thin lines denote the average genome size, gene counts, and the population sizes of individual simulations. Thick lines are the averages across simulations. The bottom rows for all subfigures shows the prevalence of TEs in individual cells on the grid over time, with colours ranging on a log scale from zero (blue/purple) to 100 (yellow).

**Supplementary Figure 4.**
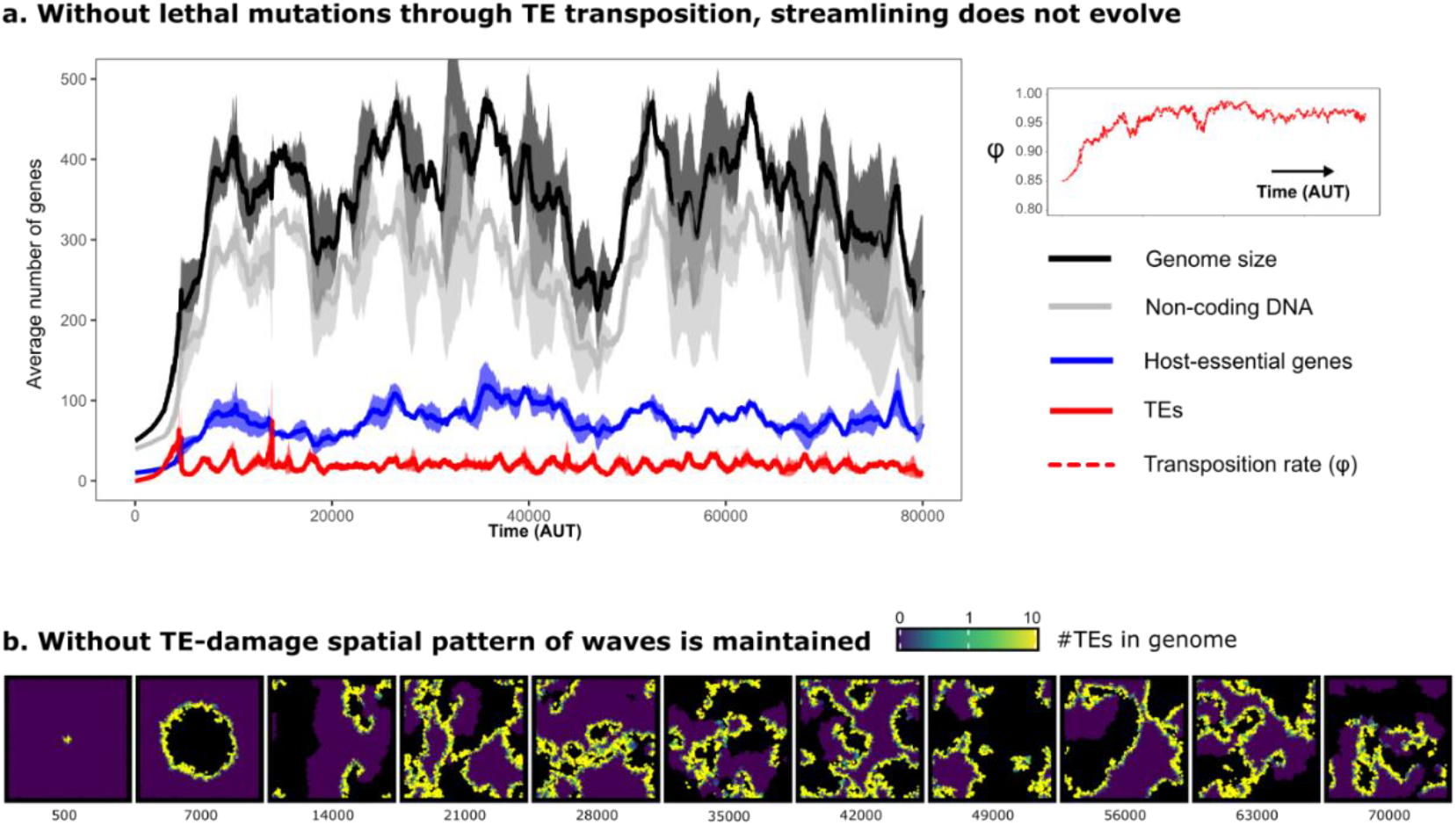
Without TE-induced damage, no streamlining evolves. However, the system still persists by forming travelling waves. **a.** The average number of genes and genome size are plotted over time. Lines show the average of five independent simulations. Shaded areas denote the standard error across simulations. Host-essential genes are shown in blue, TEs are shown in red, and non-coding positions are shown in grey. **b.** The prevalence of TEs in individual cells on the grid over time, with colours ranging on a log scale from zero (blue/purple) to 10 (yellow). All parameters are the same as **Figure 3** from the main text, but TEs do not cause inactivation of host-essential genes, and can therefore not be lethal to individual cells.

**Supplementary Figure 5.**
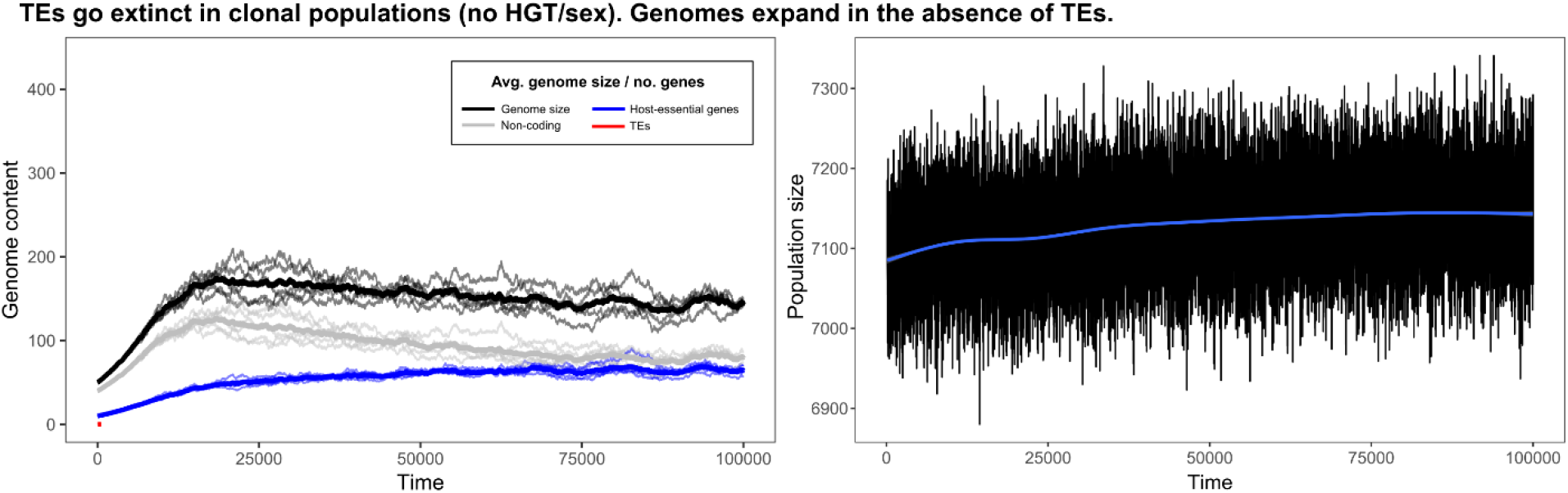
In a clonal population (no HGT/no sex) TEs immediately go extinct. In the absence of TEs, genomes expand to improve mutational robustness. For five replicate populations without DNA uptake or sex, the genome content evolution is shown. The thick lines show the average of the five lines. The graph shows that in the absence of TEs, genomes expand as the population size increases. Because there is no differential fitness (only TEs have a fitness-cost) and no differential death (only the base death rate applies), this increase in the population size can only be explained by a higher replication fidelity. Thus, the expansion of the genome (and the minor streamlining that follows!) show selection for mutational robustness.

**Supplementary Figure 6.**
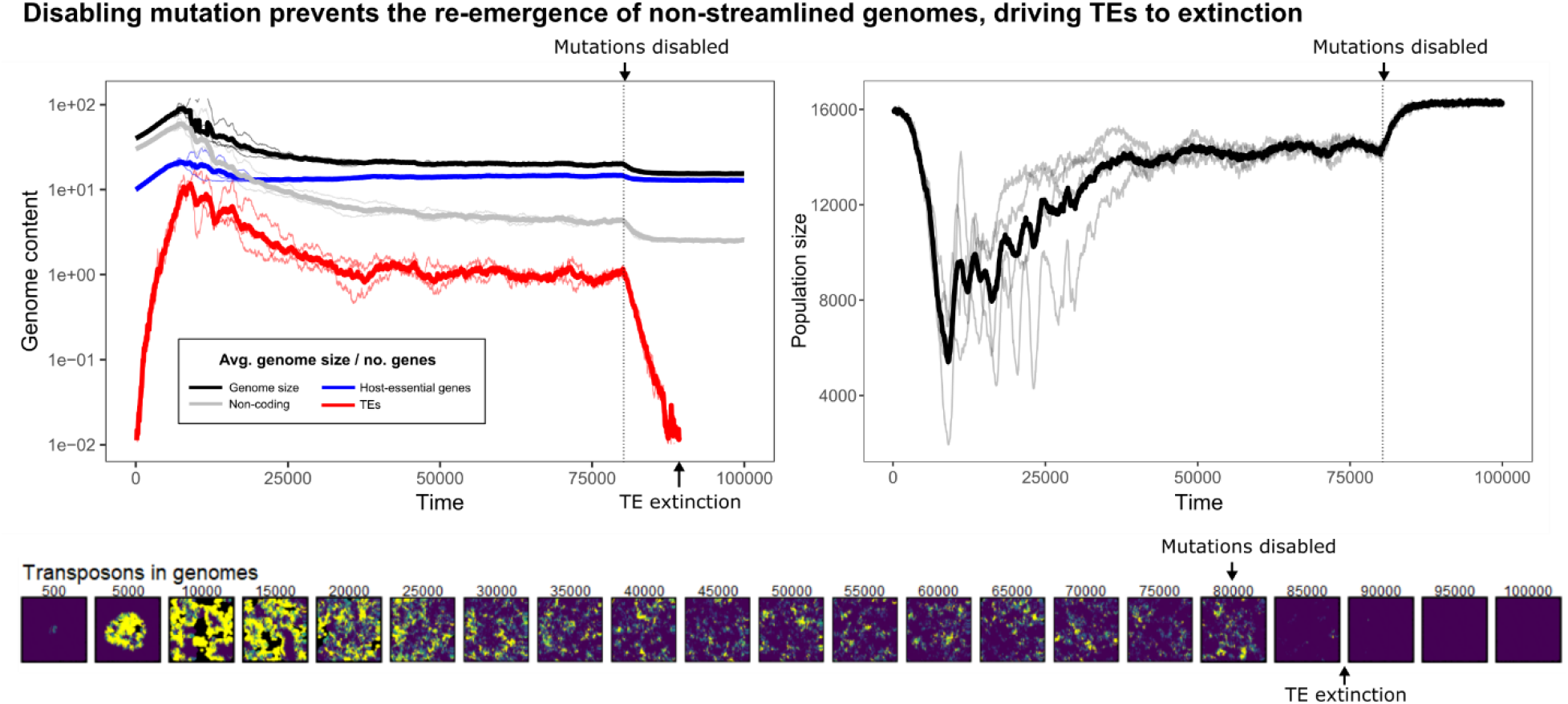
Disabling mutation after streamlining evolved breaks the rock-papers-scissors cycle (non-streamlined genomes no longer emerge), resulting in the extinction of TEs. For five replicate populations, the genome content and population size are shown. The thin lines show the average of cells in the population (for the plot on the left-hand side). The thick lines show the average of the five lines. The bottom row shows the abundance of TEs in individual cells (genomes), with colours scaling from zero (blue) to one (green) to 10 (yellow). The data shows that the rock-paper-scissors dynamic discussed in the main text depends on mutations generating non-streamlined genomes, which happens in the (local) absence of TEs. Without mutations (after T=80,000), TEs slowly go extinct as new vulnerable (non-streamlined) genomes no longer appear.

**Supplementary Figure 7.**
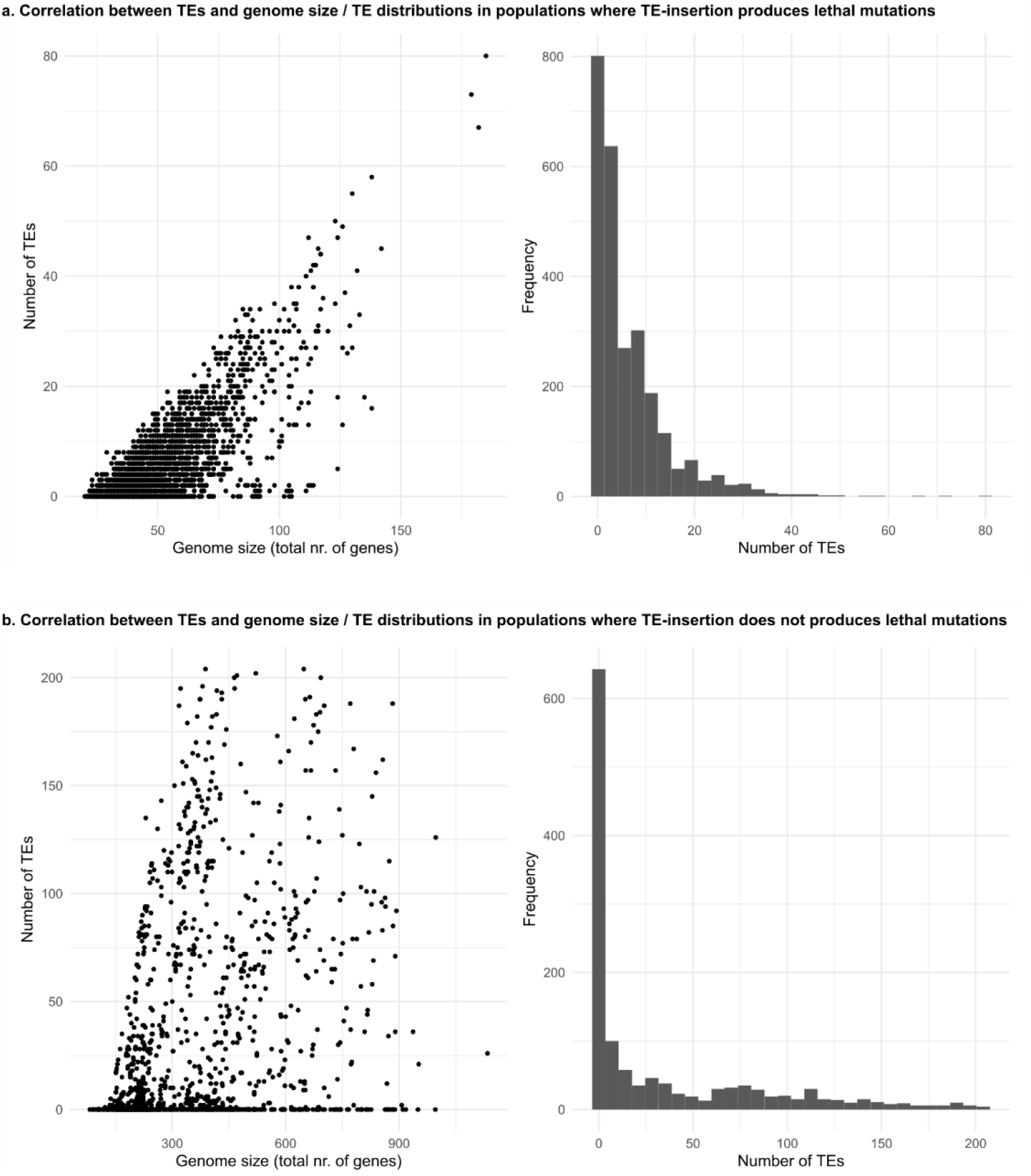
Patterns of TEs and genomes size with/without lethal mutations upon TE insertion. At the end of evolutionary simulations, all cells from the population were collected and their TE-abundance and genome size extracted. Plotting these numbers shows that populations where TEs cause harmful effects display a clear relationship of TE-abundance with genome size (a), which is much less clear in populations where TEs do not cause harmful effects. Also the distributions of TE-abundances show marked differences.

### Supplementary Video

To better visualise the process by which genomes become streamlined, a supplementary video was made. The video contains three grids which visualise different variables of the same simulation: **i)** genome size of individual genomes, **ii)** the number of transposons in genomes, and **iii)** the number of transposons in the environment. Subtitles and cartoons are included to help the viewer better understand what can be observed from these grids.

**Supplementary video 1:**
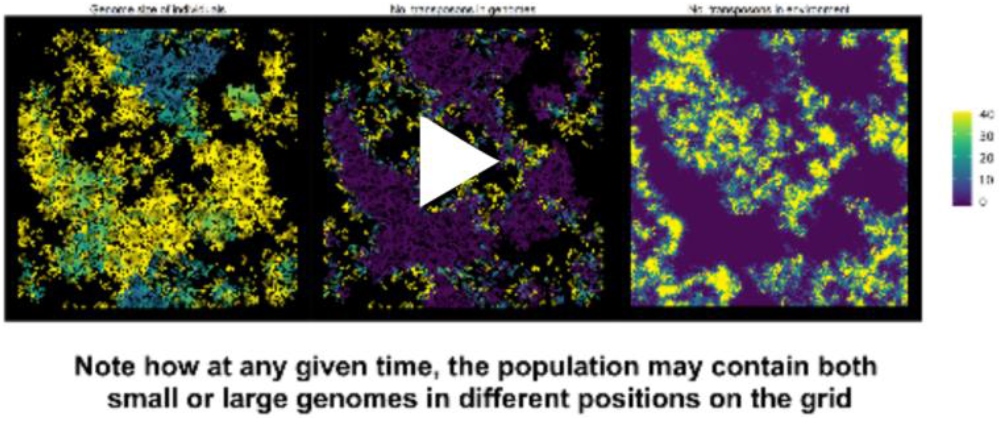
snapshot of the supplementary video available here: https://youtu.be/9KXQaoy8X0o

